# Loss of β-arrestin2 in D2 cells alters neuronal excitability in the nucleus accumbens and behavioral responses to psychostimulants and opioids

**DOI:** 10.1101/685719

**Authors:** Kirsten A. Porter-Stransky, Alyssa K. Petko, Saumya L. Karne, L. Cameron Liles, Nikhil M. Urs, Marc G. Caron, Carlos A. Paladini, David Weinshenker

## Abstract

Psychostimulants and opioids increase dopamine (DA) neurotransmission, activating D1 and D2 G protein-coupled receptors. β-arrestin2 (βarr2) desensitizes and internalizes these receptors and initiates G protein-independent signaling. Previous work revealed that mice with a global or cell-specific knockout of βarr2 have altered responses to certain drugs; however, the effects of βarr2 on the excitability of medium spiny neurons (MSNs) and its role in mediating the rewarding effects of drugs of abuse are unknown. D1-Cre and D2-Cre transgenic mice were crossed with floxed βarr2 mice to eliminate βarr2 specifically in cells containing either D1 (D1^βarr2-KO^) or D2 (D2^βarr2-KO^) receptors. We used slice electrophysiology to characterize the role of βarr2 in modulating D1 and D2 nucleus accumbens MSN intrinsic excitability in response to DA and tested the locomotor-activating and rewarding effects of cocaine and morphine in these mice. We found that eliminating βarr2 attenuated the ability of DA to inhibit D2-MSNs but had little effect on the DA response of D1-MSNs. While D1^βarr2-KO^ mice had mostly normal drug responses, D2^βarr2-KO^ mice showed dose-dependent reductions in acute locomotor responses to cocaine and morphine, attenuated locomotor sensitization to cocaine, and blunted cocaine reward measured with conditioned place preference. Both D2^βarr2-KO^ and D1^βarr2-KO^ mice displayed an enhanced conditioned place preference for the highest dose of morphine. These results indicate that D2-derived βarr2 functionally contributes to the ability of DA to inhibit D2-MSNs and multiple behavioral responses to psychostimulants and opioids, while loss of βarr2 in D1 neurons has little impact on D1-MSN excitability or drug-induced behaviors.

## Introduction

Substance abuse continues to be a public health crisis, and an urgent need exists to better understand the neurobiological effects of addictive drugs and to develop new pharmacological interventions. G protein-coupled receptors (GPCRs) are crucially involved in mediating the behavioral effects of drugs with abuse potential. Most addictive drugs increase dopamine (DA) neurotransmission in the nucleus accumbens (NAc), which stimulates D1-like and D2-like GPCRs on medium spiny neurons (MSNs) (Di Chiara & Imperato, 1988; Pierce & Kumaresan, 2006). Some drugs of abuse, such as opioids and cannabinoids, directly bind to GPCRs. β-arrestins (βarr) are important for desensitizing and internalizing GPCRs and can initiate G protein-independent signaling (Smith & Rajagopal, 2016), making them a promising target for regulating drug-induced GPCR signaling.

Accumulating evidence implicates β-arrestin2 (βarr2, also known as arrestin-3) in modulating the effects of addictive drugs (Porter-Stransky & Weinshenker, 2017). βarr2 knockout (βarr2-KO) mice, which completely lack βarr2, have blunted morphine-induced locomotion yet show an enhanced morphine conditioned place preference (CPP) (Bohn, Gainetdinov, Sotnikova et al., 2003). Conventional and neuron-selective βarr2-KOs also have reduced locomotor responses to amphetamine, but complete absence of βarr2 does not impair cocaine CPP (Beaulieu, Sotnikova, Marion et al., 2005; Bohn, Gainetdinov, Sotnikova et al., 2003; Urs, Gee, Pack et al., 2016; Zurkovsky, Sedaghat, Ahmed et al., 2017).

Alterations in βarr2 in the NAc affect cocaine-induced locomotion (Gaval-Cruz, Goertz, Puttick et al., 2014). Although the modulation of drug responses by βarr2 are thought to occur within mesolimbic circuitry, the striatal cell-types have not been fully identified. This is an important question because D1- and D2-MSNs in the NAc can have different projections and elicit distinct behavioral effects (Beaulieu & Gainetdinov, 2011; Floresco, 2015; Hikida, Morita & Macpherson, 2016; Kupchik, Brown, Heinsbroek et al., 2015; Pardo-Garcia, Garcia-Keller, Penaloza et al., 2019; Porter-Stransky, Seiler, Day et al., 2013). Inhibition of D1 transmission or a βarr2/phospho-ERK signaling complex attenuates the locomotor-activating effects of morphine (Urs, Daigle & Caron, 2011), while eliminating βarr2 in both D1- and D2-containing cells or just in D2 neurons reduces amphetamine-induced locomotion (Urs, Gee, Pack et al., 2016). Research using engineered variants of D2 receptors indicates that βarr2, independent of G protein signaling downstream of D2 receptors, contributes to the locomotor-activating effects of psychostimulants (Peterson, Pack, Wilkins et al., 2015; Rose, Pack, Peterson et al., 2018).

While mesolimbic βarr2 is implicated in the effects of drugs of abuse, there is little information regarding how βarr2 regulates the activity of D1- and D2-MSNs, and the specific cell types that mediate the impact of βarr2 on drug-induced locomotion and reward remain unclear (Gaval-Cruz, Goertz, Puttick et al., 2014; Rose, Pack, Peterson et al., 2018; Urs, Daigle & Caron, 2011; Urs, Gee, Pack et al., 2016). In the present study, we used conditional βarr2-KO mice that lack βarr2 specifically in D1 or D2 receptor-containing cells (D1^βarr2-KO^ and D2^βarr2-KO^) (Urs, Gee, Pack et al., 2016) to investigate the contribution of endogenous βarr2 to (1) DA-mediated changes in D1- and D2-MSN excitability, and (2) the locomotor-activating and rewarding effects of the psychostimulant cocaine and opioid morphine.

## Materials and Methods

### Subjects

To generate mice lacking βarr2 in either D1- or D2-containing cells, C57BL6/J D1-Cre and D2-Cre mice (Mutant Mouse Regional Resource Center, strain #036916-UCD and 017263-UCD, respectively) were crossed onto a homozygous floxed βarr2 background (Urs, Gee, Pack et al., 2016). D1^βarr2-KO^, D2^βarr2-KO^, and homozygous floxed βarr2 lacking Cre (control) mice were used for all behavioral experiments. For electrophysiology and immunohistochemisty experiments, D1^βarr2-KO^ and D2^βarr2-KO^ mice were crossed with floxed tdTomato mice to facilitate the visualization of D1 (D1^βarr2-KO/tdTomato^) and D2 (D2 ^βarr2-KO/tdTomato^) MSNs, respectively. D1-tdTomato mice (obtained from The Jackson Laboratory, stock #016204) and D2-GFP mice (obtained from Mutant Mouse Regional Resource Center, strain #036931-UCD) with normal βarr2 levels were used as controls in electrophysiology experiments. All strains were maintained on a C57Bl/6 background. Genotypes were confirmed by PCR. Approximately equal numbers of male and female mice of each genotype were included in experiments, and subjects were kept on a 12:12 light/dark cycle with lights on at 7:00 AM. Behavioral testing occurred during the light cycle of 3-6 months old mice. All procedures were approved by the Institutional Animal Care and Use Committee (IACUC) of Emory University and the University of Texas-San Antonio. To confirm βarr reductions in the appropriate cells of the conditional KOs, tissue was prepared and immunohistochemistry was performed, as detailed in the Supplementary Materials and Methods and previously described (Porter-Stransky, Centanni, Karne et al., 2019).

### Slice electrophysiology

Electrophysiological recordings of NAc MSNs from D1^βarr2-KO/tdTomato^, D2^βarr2-KO/tdTomato^, D1-tdTomato, and D2-GFP mice were conducted as described in the Supplementary Materials and Methods.

### Drugs

Cocaine and morphine were supplied by the National Institute on Drug Abuse Drug Supply Program. Doses were chosen based upon previous studies and to capture potential dose-dependent effects (Bohn, Gainetdinov, Sotnikova et al., 2003; Gaval-Cruz, Goertz, Puttick et al., 2014; Schank, Ventura, Puglisi-Allegra et al., 2006). Drugs were mixed in 0.9% bacteriostatic saline and injected at a volume of 10 ml/kg. DA was purchased from Tocris (Minneapolis, MN).

### Drug-induced locomotion

Locomotor testing was conducted in San Diego Instruments (La Jolla, CA) locomotor chambers as described in the Supplementary Materials and Methods.

### CPP

A 5-day paradigm in a 3-chamber CPP apparatus (San Diego Instruments) was used as previously described (Schank, Ventura, Puglisi-Allegra et al., 2006) and detailed in the Supplementary Materials and Methods.

### Statistics

Because the data were normally distributed, parametric tests were used to analyze the data in SPSS Version 25 and GraphPad Prism Version 7. An alpha level of 0.05 was used to determine statistical significance. Mixed model ANOVAs (genotype as a between-subjects factor and DA application as a repeated-measures factor) were used to analyze DA-induced changes in maximum number of spikes, slope of F-I curves, half maximal current, and rheobase in electrophysiology experiments. Unpaired t-tests were used to compare the baseline data between the conditional βarr2-KO cells and control cells.

For locomotor experiments, the 90-min habituation session prior to drug injection was analyzed separately from the 2-h drug-induced locomotion post injection. For acute drug-induced locomotor experiments, ambulations in 5-minute time bins were analyzed with a linear mixed model because of its ability to handle repeated measures data in which observations are not independent (Aragona, Cleaveland, Stuber et al., 2008; Vander Weele, Porter-Stransky, Mabrouk et al., 2014). D1^βarr2-KO^ and D2 ^βarr2-KO^ mice were compared to controls. A mixed model ANOVA was used to analyzed the first 30 minutes (binned into 1 data point) of drug-induced locomotion followed by Dunnett’s multiple comparisons test. To analyzed locomotor sensitization to cocaine, a mixed model ANOVA also was used comparing cocaine-induced locomotion on day 1 to day 7 (genotype as between-subjects factor and session as repeated-measures factor) followed by a posthoc test with Bonferroni correction. To analyze session-by-session data in the cocaine sensitization experiment, a mixed model ANOVA (genotype as between-subjects factor and session as repeated-measures factor) and a Dunnett’s multiple comparisons test were used. An unpaired t-test was used to compare saline-induced locomotion of controls with a D1-Cre or D2-Cre parent and to compare cocaine-induced locomotion during CPP of D2 ^βarr2-KO^ mice to controls. To test whether subjects of each genotype developed a CPP for each dose of drug, mixed model ANOVAs (time as a repeated factor and dose as a between-subjects factor) and posthoc tests with Bonferroni corrections were used to compare differences in preference (time spent in saline-paired chamber subtracted from time spent in drug-paired chamber) between the pre-test (prior to conditioning) and the post-test (after conditioning). Cohen’s *d* was calculated to determine effect sizes for statistically significant effects.

Although male and female subjects were included in all experimental groups, sex was not included as a factor in analyses. Sex differences were not the focus of the present study, and the experiments were not sufficiently powered to detect them. However, we visually inspected the data based on sex and saw no apparent differences.

## Results

### Generation of mice lacking βarr2 in D1 or D2 cells

D1-Cre and D2-Cre mice were crossed onto a homozygous floxed βarr2 background to generate KOs lacking βarr2 specifically in either D1 or D2 cells (Urs, Gee, Pack et al., 2016). To visualize MSNs lacking βarr2, a floxed tdTomato allele was introduced in the conditional KOs. Thus, tdTomato should be expressed preferentially in cells lacking βarr2, while cells with normal βarr2 levels should be devoid of tdTomato. Immunohistochemistry and electrophysiology experiments were performed on slices taken from the NAc (Figure 1A-C). Consistent with previous studies, approximately half of the analyzed neurons in the NAc expressed tdTomato, indicating equivalent percentages of D1- and D2-MSNs in both conditional βarr2-KOs (Figure 1D). Whereas βarr2 immunoflorescence was observed in the majority (∼75%) of non-tdTomato cells (primarily D2 cells in D1^βarr2-KO^ mice and D1 cells in D2^βarr2-KO^ mice; Figure 1E), βarr2 immunoflorescence only was detected in a minority (∼25%) of tdTomato cells (D1 cells in D1^βarr2-KO^ mice and D2 cells in D2^βarr2-KO^ mice; Figure 1F). Therefore, βarr2 was mostly absent from D1-MSNs but not D2-MSNs in D1^βarr2-KO^ mice, while the opposite was true in D2^βarr2-KO^ mice. The observed βarr2 signal was not due to non-specific binding of the secondary antibody (Figure S1).

**Figure 1.**
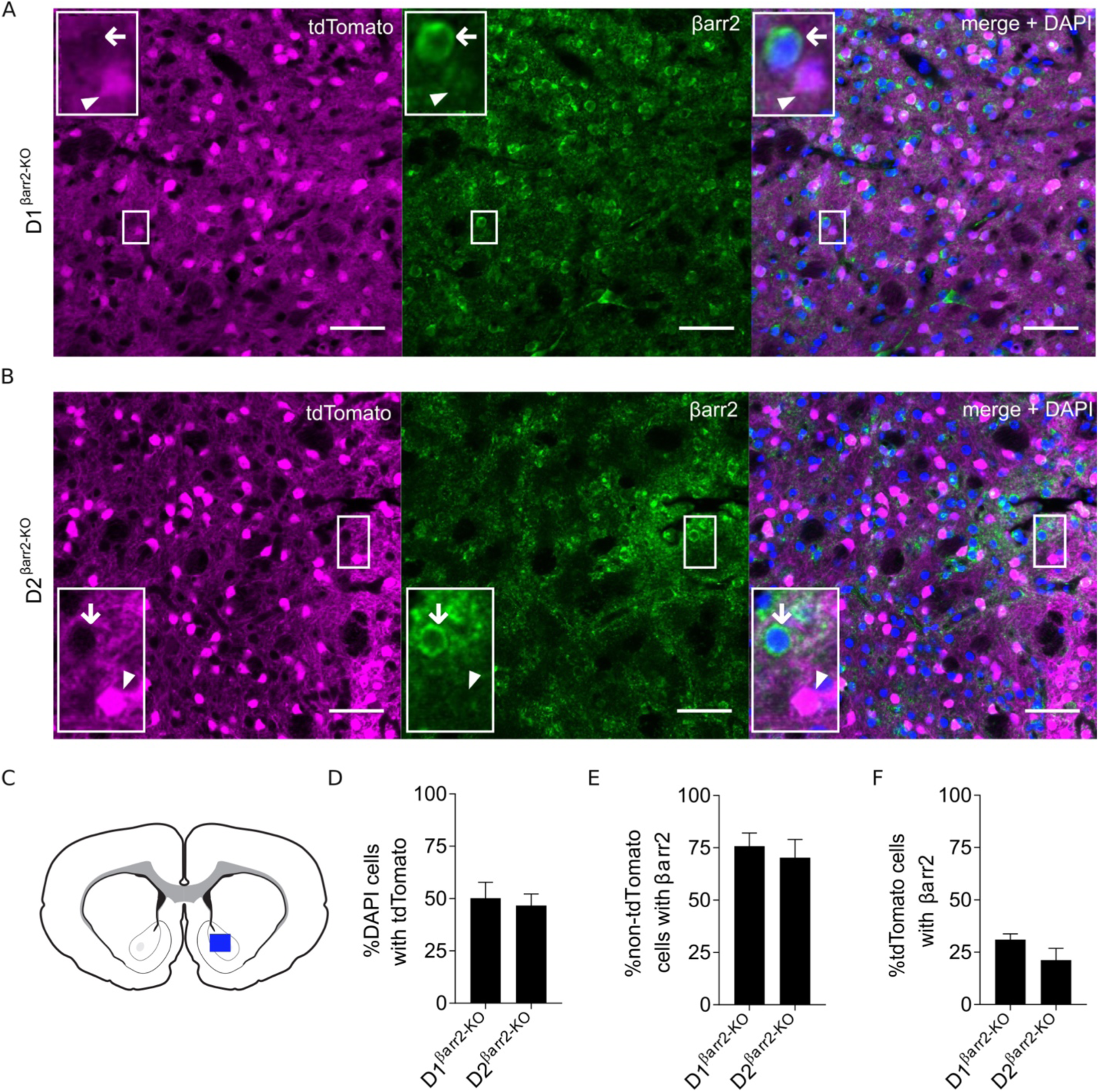
Immunohistochemistry of conditional βarr2-KOs. **A-B)** Representative images of endogenous tdTomato fluorescence of D1- (A) or D2- (B) MSNs (magenta) from D1^βarr2-KO/tdTomato^ and D2 ^βarr2-KO/tdTomato^ mice, respectively. Green indicates βarr2 immunofluorescence. Right panel shows merge of tdTomato, βarr2, and DAPI (blue). Arrowheads indicate examples of D1-MSNs (A) or D2-MSNs (B) not containing βarr2. Arrows point to non-tdTomato cells containing βarr2. All scale bars = 50 µm. **C-F)** Images were taken from the NAc (blue rectangle; C) and quantification of DAPI, tdTomato, and βarr2 in NAc cells was performed. Approximately half the analyzed cells in each genotype expressed tdTomato (indicating D1- or D2-MSNs; D). βarr2 co-localized in most non-tdTomato-expressing cells (E) but was absent in the majority of tdTomato-expressing cells (F). Error bars represent SEM; n = 2-3 mice per group.

### βarr2 preferentially modulates DA-induced activity in D2-MSNs

To determine the contribution of βarr2 to DA-induced changes in NAc D1- and D2-MSN excitability, we performed ex vivo whole cell recordings. No statistically significant differences were observed for baseline electrophysiological properties between D1^βarr2-KO^ and control D1-MSNs (Figure 2; maximum firing rate, *t*_(31)_ = 1.903, *p* = 0.066; slope of F-I curve, *t*_(31)_ = 0.945, *p* = 0.352; half maximum current, *t*_(31)_ = 0.922, *p* = 0.364; rheobase, *t*_(31)_ = 0.661, *p* = 0.514) or between D2 ^βarr2-KO^ and control D2-MSNs (Figure 3; maximum firing rate, *t*_(40)_ = 0.380, *p* = 0.706; slope of F-I curve, *t*_(40)_ = 0.466, *p* = 0.644; half maximum current, *t*_(40)_ = 1.517, *p* = 0.137; rheobase, *t*_(40)_ = 1.766, *p* = 0.085).

**Figure 2.**
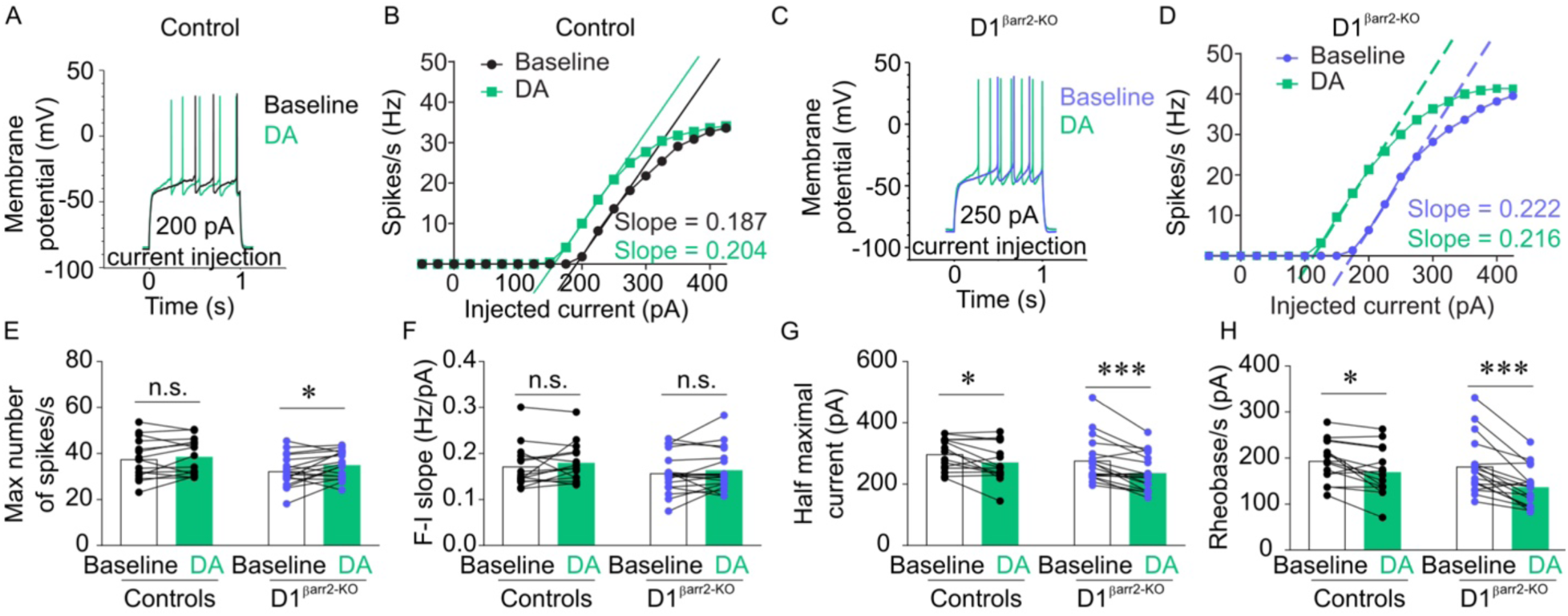
Effects of DA on NAc D1-MSNs lacking βarr2. **A-D)** Representative, single-cell traces (A, C) and F-I curves (B, D) of NAc D1-MSN activity at indicated levels of current injection in response to bath application of DA (60 µM) from control and D1^βarr2-KO^ mice. **E-H)** DA increased the maximum firing rate of D1-MSNs in D1^βarr2-KO^ but not control mice (E). DA had similar effects on F-I slope (F), half maximal current (G), and rheobase (H) in D1^βarr2-KO^ and control D1-MSNs. \**p* < 0.05, \*\**p* < 0.001, n.s. = not statistically significant, error bars represent SEM, n = 15-18 cells per genotype.

**Figure 3.**
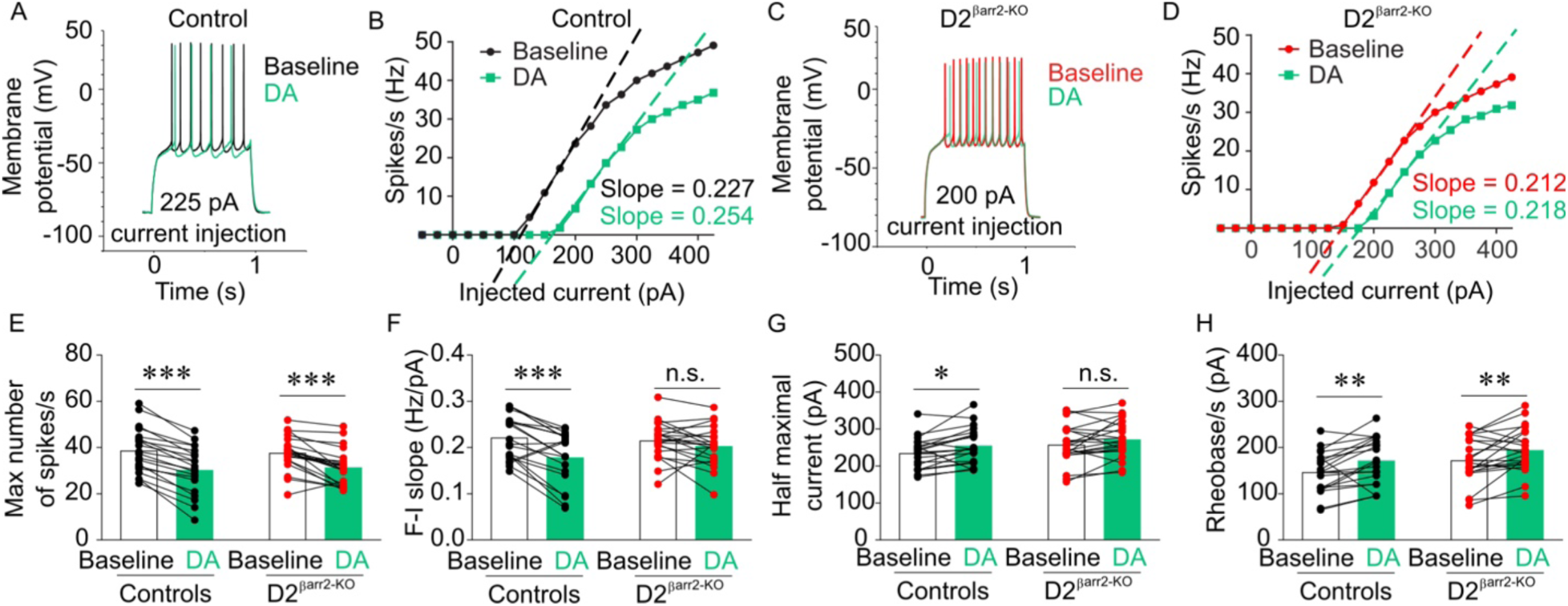
Effects of DA on NAc D2-MSNs lacking βarr2. **A-D)** Representative, single-cell examples of NAc D2-MSN responses from control and D2^βarr2-KO^ mice. The example traces (A, C) are representative of the typical changes observed in response to DA (60 µM), and the example F-I curves show single-cell responses to DA at indicated levels of current injection (B, D) to highlight the changes in slope of the F-I curves from control but not D2^βarr2-KO^ cells. **E-H)** DA decreased the maximum number of spikes in control and D2-MSNs lacking βarr2 (E). DA reduced the slope of the F-I curve (F) and increased the half maximal current (G) in control D2-MSNs but not in D2-MSNs lacking βarr2. DA increased rheobase in both control and D2^βarr2-KO^ D2-MSNs. \**p* < 0.05, \*\**p* < 0.01, \*\*\**p* < 0.001, n.s. = not statistically significant, error bars represent SEM, n = 20-22 cells per genotype.

When DA (60 µM) was bath-applied to NAc slices, an increase in intrinsic excitability in D1 neurons was observed in both D1^βarr2-KO/tdTomato^ and D1^tdTomato^ mice (Figure 2A-D). DA increased the maximum firing rate of D1 cells lacking βarr2; however, DA did not significantly alter the maximum firing rate of D1-MSNs in control mice (Figure 2E; main effect of DA *F*_(1,31)_ = 5.114, *p* = 0.031; interaction *F*_(1,31)_ = 0.958, *p* = 0.335; controls, *p* = 0.392; D1^βarr2-KO^, *p* = 0.022, Cohen’s *d* = 0.526). DA did not significantly alter the slope of F-I curves, regardless of βarr2 content (Figure 2F; main effect of DA, *F*_(1,31)_ = 2.042, *p* = 0.163; interaction *F*_(1,31)_ = 0.014, *p* = 0.907; controls, *p* = 0.303; D1^βarr2-KO^, *p* = 0.338). DA caused a significant decrease in the half maximum current (Figure 2G; main effect of DA *F*_(1,31)_ = 25.770, *p* < 0.001; interaction *F*_(1,31)_ = 1.089, *p* = 0.305; controls, *p* = 0.010, Cohen’s *d* = 0.697; D1^βarr2-KO^, *p* < 0.001, Cohen’s *d* = 1.078) and the threshold current necessary to generate an action potential (rheobase, Figure 2H; main effect of DA *F*_(1,31)_ = 32.513, *p* < 0.001; interaction *F*_(1,31)_ = 2.830, *p* = 0.103; controls, *p* = 0.011 Cohen’s *d* = 0.809; D1^βarr2-KO^, *p* < 0.001, Cohen’s *d* = 1.177) in both D1^βarr2-KO^-MSNs and control D1-MSNs. Together, these data indicate that βarr2 only modestly affected a single aspect of D1-MSNs’ responses to DA, as indicated by an increase in the maximum number of spikes when the D1 neurons lack βarr2, while all other measures were comparable in D1^βarr2-KO^ and control D1-MSNs.

By contrast, there were several marked genotype differences in the D2-MSN DA response. DA (60 µM) inhibited D2-MSN excitability in both D2^βarr2-KO/tdTomato^ and D2-GFP mice, but the effect was much more profound in D2-MSNs with normal βarr2 content (Figure 3A-D). DA also significantly decreased maximum firing rate in control D2-MSNs and D2-MSNs lacking βarr2 (Figure 3E; main effect of DA *F*_(1,40)_ = 97.884, *p* < 0.001; interaction *F*_(1,40)_ = 2.047, *p* = 0.160; controls, *p* < 0.001, Cohen’s *d* = 2.083; D2^βarr2-KO^, *p* < 0.001, Cohen’s *d* = 1.162). However, DA caused a significant decrease in the slope of the F-I curve in control D2-MSNs but not in D2-MSNs lacking βarr2 (Figure 3F; main effect of DA *F*_(1,40)_ = 20.991, *p* < 0.001; interaction *F*_(1,40)_ = 6.726, *p* = 0.013; controls, *p* < 0.001, Cohen’s *d* = 1.037; D2^βarr2-KO^, *p* = 0.157). Similarly, DA significantly increased the half maximal current required for an action potential in control D2-MSNs but its effects on this parameter in D2^βarr2-KO^-MSNs did not reach statistical significance (Figure 3G; main effect of DA *F*_(1,40)_ = 10.772, *p* = 0.002; interaction *F*_(1,40)_ = 0.329, *p* = 0.569; controls, *p* = 0.011, Cohen’s *d* = 0.649; D2^βarr2-KO^, *p* = 0.057). DA increased rheobase in both control and D2-MSNs lacking βarr2 (main effect of DA *F*_(1,40)_ = 18.427, *p* < 0.001; interaction *F*_(1,40)_ = 0.060, *p* = 0.808; controls, *p* = 0.003, Cohen’s *d* = 0.710; D2^βarr2-KO^, *p* = 0.006, Cohen’s *d* = 0.618). Together, these results indicate that βarr2 is required for full D2-MSN responses to DA; eliminating βarr2 in D2-MSNs makes these neurons less sensitive to the inhibitory effects of DA as evidenced by blunted inhibition and lack of DA-induced reduction in the slope of F-I curves and half maximal current.

### Loss of βarr2 in D2 but not D1 cells attenuates the locomotor-activating effects of cocaine and morphine

Mice lacking βarr2 in all neurons have attenuated locomotor responses to amphetamine and morphine (Beaulieu, Sotnikova, Marion et al., 2005; Bohn, Gainetdinov, Sotnikova et al., 2003; Urs, Gee, Pack et al., 2016), which are DA-dependent behaviors, and we identified greater alterations in the cellular responses of βarr2-lacking D2-MSNs to DA compared to D1-MSNs. D1^βarr2-KO^, D2^βarr2-KO^, and control mice exhibited nearly identical levels of novelty-induced locomotion (Figure S2A; D1^βarr2-KO^ main effect of genotype *F*_(1,20.222)_ = 0.534, *p* = 0.473, interaction *F*_(17,318.637)_ = 0.692, *p* = 0.811; D2^βarr2-KO^ main effect of genotype *F*_(1,22.847)_ = 0.588, *p* = 0.451, interaction *F*_(17,302.920)_ = 0.678, *p* = 0.824) and saline-induced locomotion (Figure S2B; D1^βarr2-KO^ main effect of genotype *F*_(1,60.172)_ = 1.046, *p* = 0.310, interaction *F*_(24,306.375)_ = 1.076, *p* = 0.370; D2^βarr2-KO^ main effect of genotype *F*_(1,76.874)_ = 0.927, *p* = 0.339, interaction *F*_(24,294.475)_ = 0.505, *p* = 0.976), indicating that βarr2 in these cells is not required for normal locomotion in the absence of drug. Furthermore, locomotor responses to a saline injection did not differ regardless of whether control mice came from a D1-Cre or D2-Cre litter (Figure S2C; *t*_(26)_ = 0.371, *p* = 0.714).

To determine whether βarr2 in D1 or D2 cells modulates the acute locomotor-activating effects of drugs of abuse, ambulations following administration of morphine or cocaine were examined. Cocaine-induced locomotion in D1^βarr2-KO^ mice was comparable to controls at all doses tested (Figure 4A-C; 5 mg/kg main effect of genotype *F*_(1,51.878)_ = 0.856, *p* = 0.359, interaction *F*_(24,322.681)_ = 1.109, *p* = 0.332; 10 mg/kg main effect of genotype *F*_(1,45.393)_ = 0.400, *p* = 0.530, interaction *F*_(24,342.980)_ = 1.674, *p* = 0.026; 20 mg/kg main effect of genotype *F*_(1,23.598)_ = 0.023, *p* = 0.880, interaction *F*_(24,418.171)_ = 1.034, *p* = 0.420) but was significantly reduced in D2^βarr2-KO^ mice at the 10 mg/kg dose (Figure 4A-C; 5 mg/kg main effect of genotype *F*_(1,64.691)_ = 0.749, *p* = 0.390, interaction *F*_(24,305.341)_ = 1.201, *p* = 0.239; 10 mg/kg main effect of genotype *F*_(1,50.427)_ = 6.739, *p* = 0.012, interaction *F*_(24,335.365)_ = 1.770, *p* = 0.016; 20 mg/kg main effect of genotype *F*_(1,24.874)_ = 0.145, *p* = 0.707, interaction *F*_(24,406.685)_ = 1.122, *p* = 0.315). We later replicated this finding by examining ambulations in a separate group of mice during CPP training; D2 ^βarr2-KO^ mice administered 10 mg/kg cocaine once again exhibited blunted cocaine-induced locomotion compared to controls (Figure 4D; *t*_(14)_ = 2.188, *p* = 0.046, Cohen’s *d* = 3.095).

**Figure 4.**
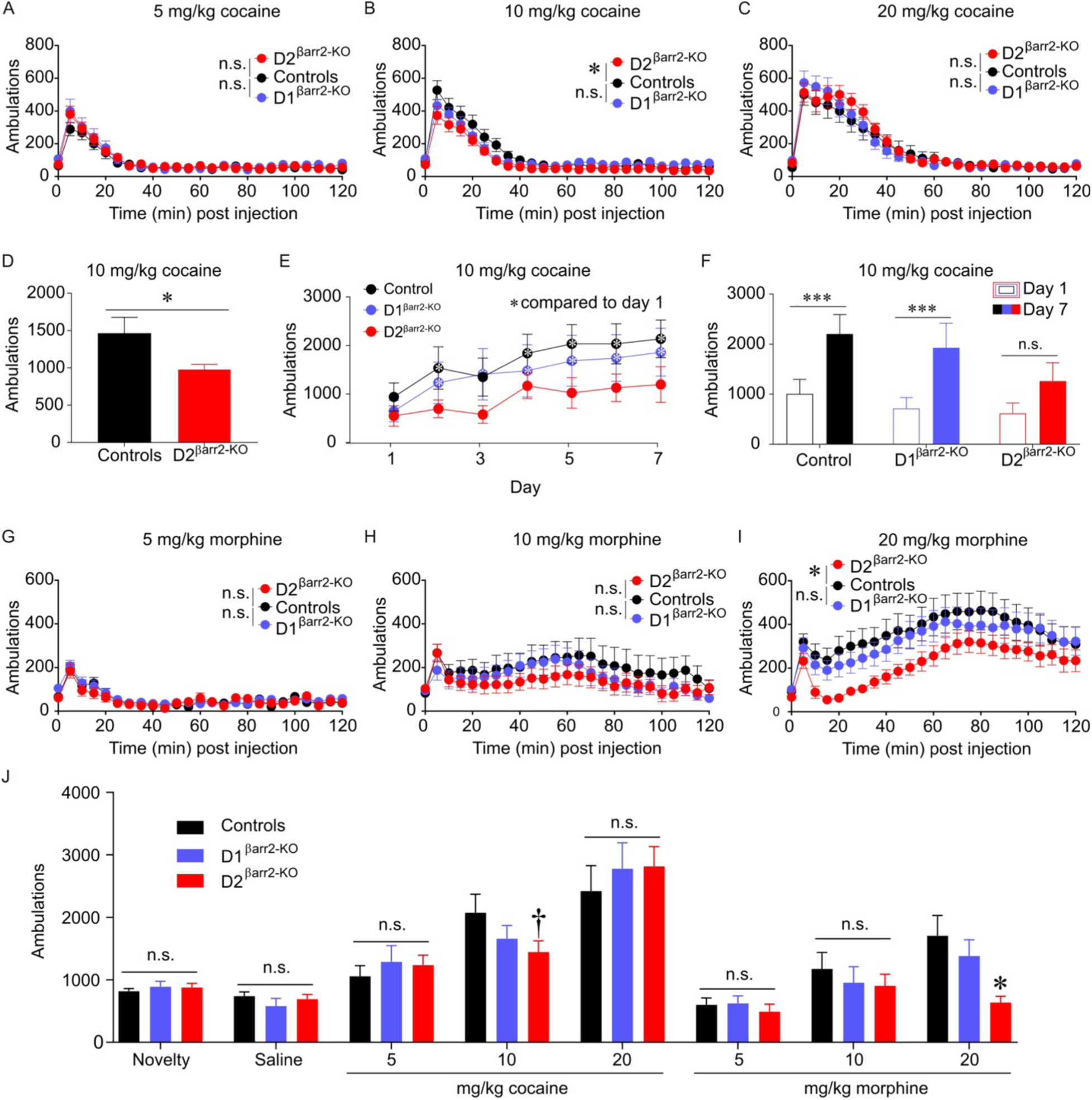
Loss of βarr2 in D2 cells attenuates the acute locomotor-activating effects of cocaine and morphine dose-dependently. **A-D)** Cocaine-induced locomotion was dose-dependently blunted in D2^βarr2-KO^ mice but not D1 ^βarr2-KO^ mice (A, 5 mg/kg cocaine; B, 10 mg/kg cocaine; C, 20 mg/kg cocaine; \**p* < 0.05). The reduced locomotor activation of D2^βarr2-KO^ mice was replicated in a separate group of subjects (D, 10 mg/kg cocaine; \**p* < 0.05). **E-F)** In a sensitization experiment, cocaine (10 mg/kg)-induced locomotion was measured for 7 consecutive days. Control and D1^βarr2-KO^ mice showed elevated cocaine-induced locomotion on on several days compared to day 1, whereas D2^βarr2-KO^ mice did not (E, \**p* < 0.05 compared to day 1). Control and D1^βarr2-KO^ mice sensitized to cocaine by the end of the experiment (day 7 compared to day 1, \*\*\**p* < 0.001), but D2^βarr2-KOs^ did not have significantly elevated cocaine-induced locomotion on day 7 compared to day 1 (F). **G-I)** Morphine-induced locomotion was dose-dependently attenuated in D2^βarr2-KO^ mice but not D1 ^βarr2-KO^ mice (G, 5 mg/kg morphine; H, 10 mg/kg morphine; H, 20 mg/kg morphine; \**p* < 0.05). **G)** Acute drug-induced locomotor data showing cumulative ambulations 30 minutes post-injection (\**p* < 0.05; †*p* = 0.08). n.s. = not statistically significant, error bars represent SEM, n = 8-12 mice per group.

Because βarr2 is necessary for amphetamine sensitization (Zurkovsky, Sedaghat, Ahmed et al., 2017), we tested whether βarr2 also contributes to cocaine locomotor sensitization. Subjects received cocaine (10 mg/kg, i.p.) daily for 7 days. Analysis of cocaine-induced activity by day revealed that control and D1^βarr2-KO^ mice displayed significantly greater locomotor responses to cocaine in subsequent sessions compared to day 1, but D2^βarr2-KO^ mice did not (Figure 4E; main effect of time *F*_(6,174)_ = 15.518, *p* < 0.001; main effect of genotype *F*_(2,29)_ = 0.943, *p* = 0.943; interaction *F*_(12,174)_ = 0.762, *p* = 0.688; posthoc test with Dunnett’s correction revealed days 2, 4-7 compared to day 1 *p* < 0.05 in controls; days 2-7 compared to day 1 *p* < 0.05 in D1^βarr2-KO^ mice; none of days 2-7 were significantly higher than day 1, *p* > 0.05, in D2^βarr2-KO^ mice). Indeed, whereas control and D1^βarr2-KO^ mice sensitized to cocaine, D2 ^βarr2-KO^ mice did not exhibit a significantly enhanced response to cocaine on day 7 compared to day 1 (Figure 4F; main effect of time *F*_(1,29)_ = 42.524, *p* < 0.001; interaction *F*_(2,29)_ = 1.241, *p* = 0.304; posthoc test with Bonferroni corrections, controls *p* < 0.001, Cohen’s *d* = 1.360; D1^βarr2-KO^ *p* < 0.001, Cohen’s *d* = 1.191; D2^βarr2-KO^ *p* = 0.131).

To examine whether βarr2 is similarly important in D2 cells for opioid responses, we examined morphine-induced locomotion in the three genotypes. Morphine-induced locomotion in D1^βarr2-KO^ was similar to controls at all doses tested (Figure 4D-F; 5 mg/kg main effect of genotype *F*_(1,54.613)_ = 0.356, *p* = 0.553, interaction *F*_(24,306.834)_ = 0.902, *p* = 0.599; 10 mg/kg main effect of genotype *F*_(1,33.092)_ = 0.626, *p* = 0.434, interaction *F*_(24,442.906)_ = 1.075, *p* = 0.369; 20 mg/kg main effect of genotype *F*_(1,26.887)_ = 0.243, *p* = 0.626, interaction *F*_(24,458.327)_ = 0.510, *p* = 0.976). By contrast, morphine-induced locomotion was blunted in D2^βarr2-KO^ mice at the highest dose tested (Figure 4D-F; 5 mg/kg main effect of genotype *F*_(1,64.848)_ = 0.125, *p* = 0.725, interaction *F*_(24,300.471)_ = 1.394, *p* = 0.107; 10 mg/kg main effect of genotype *F*_(1,32.383)_ = 1.529, *p* = 0.225, interaction *F*_(24,438.530)_ = 0.749, *p* = 0.800; 20 mg/kg main effect of genotype *F*_(1,26.370)_ = 4.946, *p* = 0.035, interaction *F*_(24,453.726)_ = 1.620, *p* = 0.033).

To encapsulate the locomotor effects, the first 30 minutes of locomotor activity were summed for each condition (Figure 4J). Consistent with the results above, analysis of the binned locomotor activity data revealed that D1^βarr2-KO^ mice did not differ from controls in any tested condition, but D2^βarr2-KO^ mice exhibited reduced locomotor activity compared to controls following the high dose of morphine and a strong trend for the medium dose of cocaine (main effect of drug condition, *F*_(7,210)_ = 41.978, *p* < 0.001; interaction, *F*_(14,210)_ = 1.830, *p* = 0.036; Dunnett’s posthoc test: all conditions *p* > 0.3 except controls vs D2^βarr2-KO^ 20 mg/kg morphine *p* = 0.001, Cohen’s *d* = 1.336; 10 mg/kg cocaine *p* = 0.080, Cohen’s *d* = 0.778).

### Loss of βarr2 in D2 cells blunts the rewarding effects of cocaine but not morphine

Because conventional βarr2-KO mice have enhanced morphine CPP and normal cocaine CPP (Bohn, Gainetdinov, Sotnikova et al., 2003), we used this paradigm to determine the effects of cell type-specific βarr2 deletion on drug reward. Both control (Figure 5A; main effect of time *F*_(1,28)_ = 38.307, *p* < 0.001; interaction *F*_(3,28)_ = 3.651, *p* = 0.024; saline, *p* = 0.547; 5 mg/kg, *p* = 0.015, Cohen’s *d* = 0.891; 10 mg/kg, *p* < 0.001, Cohen’s *d* = 1.302; 20 mg/kg, *p* < 0.001, Cohen’s *d* = 2.966) and D1^βarr2-KO^ mice (Figure 5B; main effect of time *F*_(1,28)_ = 20.723, *p* < 0.001; interaction *F*_(3,28)_ = 2.066, *p* = 0.127; saline, *p* = 0.794; 5 mg/kg, *p* = 0.005, Cohen’s *d* = 0.838; 10 mg/kg, *p* = 0.031, Cohen’s *d* = 0.694; 20 mg/kg, *p* = 0.001, Cohen’s *d* = 3.093) developed a CPP for each dose of cocaine tested. By contrast, D2^βarr2-KO^ mice only showed a significant preference for the highest dose of cocaine tested (Figure 5C; main effect of time *F*_(1,28)_ = 11.182, *p* = 0.002; interaction *F*_(3,28)_ = 1.042, *p* = 0.389; saline, *p* = 0.265; 5 mg/kg, *p* = 0.246; 10 mg/kg, *p* = 0.255; 20 mg/kg, *p* = 0.003, Cohen’s *d* = 1.007). These data reveal that βarr2 in D2-, but not D1-, containing cells modulates the rewarding effects of cocaine.

**Figure 5.**
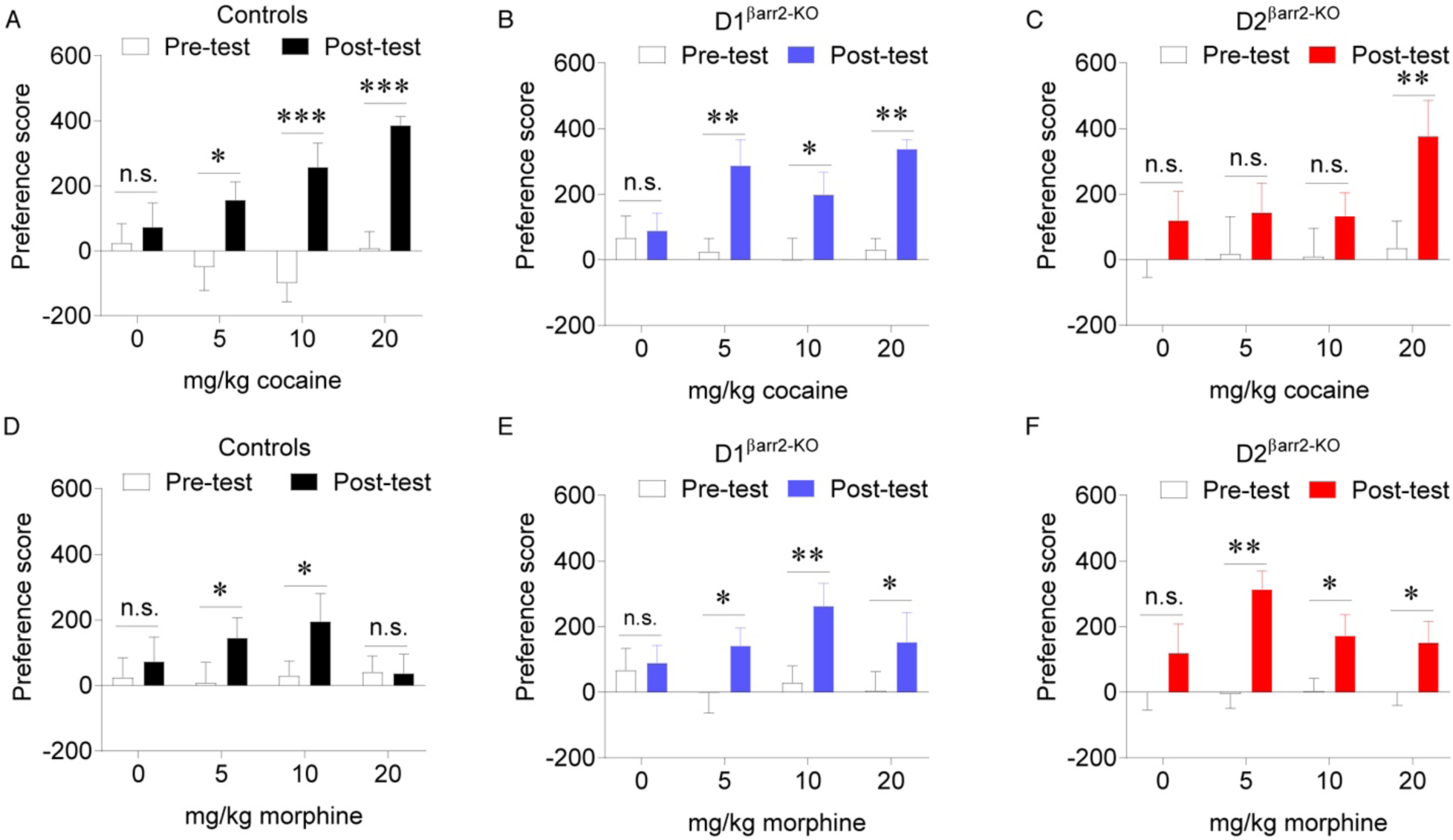
Loss of βarr2 in DA receptor-containing cells alters the rewarding effects of cocaine and morphine dose-dependently. **A-C)** For each subject’s pre- and post-test, a preference score was calculated by subtracting the time (s) spent in the saline-paired side from the time (s) spent in the drug-paired side. Whereas control (A) and D1^βarr2-KO^ (B) mice developed a CPP for each dose of cocaine tested (5, 10, 20 mg/kg), D2^βarr2-KO^ mice only developed a CPP for the highest tested dose of cocaine (C). **D-F)** Control (D) mice showed a CPP for morphine at the 5 and 10 mg/kg doses, but not 20 mg/kg dose, while both D1^βarr2-KO^ (E) and D2^βarr2-KO^ (F) mice developed a CPP for all tested doses of morphine. \**p* < 0.05, \*\**p* < 0.01, \*\*\**p* < 0.001, n.s. = not statistically significant, error bars represent SEM, n = 8-12 mice per group.

While control mice developed a CPP to the low and medium but not high dose of morphine tested (Figure 5D; main effect of time *F*_(1,34)_ = 6.093, *p* = 0.019; interaction *F*_(3,34)_ = 1.982, *p* = 0.135; saline, *p* = 0.519; 5 mg/kg, *p* = 0.040, Cohen’s *d* = 0.417; 10 mg/kg, *p* = 0.011, Cohen’s *d* = 0.454; 20 mg/kg, *p* = 0.652), all doses of morphine supported a CPP in D1^βarr2-KO^ and D2 ^βarr2-KO^ mice (Figure 5E-F; D1 ^βarr2-KO^ main effect of time *F*_(1,29)_ = 17.030, *p* < 0.001; interaction *F*_(3,29)_ = 1.652, *p* = 0.199; saline, *p* = 0.737; 5 mg/kg morphine, *p* = 0.041, Cohen’s *d* = 1.126; 10 mg/kg morphine, *p* = 0.002, Cohen’s *d* = 1.284; 20 mg/kg morphine, *p* = 0.027, Cohen’s *d* = 0.711; D2^βarr2-KO^ main effect of time *F*_(1,36)_ = 24.227, *p* < 0.001; interaction *F*_(3,36)_ = 1.130, *p* = 0.350; saline, *p* = 0.160; 5 mg/kg morphine, *p* = 0.001, Cohen’s *d* = 1.692; 10 mg/kg morphine, *p* = 0.020, Cohen’s *d* = 0.633; 20 mg/kg morphine, *p* = 0.035, Cohen’s *d* = 0.600). These results indicate that βarr2 in D1 and D2 cells is not necessary for morphine CPP and suggests that a lack of βarr2 in either cell type permits higher doses of morphine to be rewarding.

## Discussion

Using transgenic mice, electrophysiology, and behavioral assays, the present study investigated the contributions of βarr2 in NAc D1- and D2-MSN responses to DA as well as to the locomotor-activating and rewarding effects of cocaine and morphine. The results indicate that loss of βarr2 in D2-MSNs reduces the inhibitory effects of DA, attenuates the locomotor-activating effects of cocaine and morphine, blunts the rewarding properties of cocaine, and enhances the rewarding properties of morphine in a dose-dependent fashion. These results provide novel insights into the effects of βarr2 on DA-induced excitability of NAc MSNs and cell-type specificity of βarr2’s functional contributions to psychostimulant reward.

βarr2 protein levels are relatively low in the dorsal and ventral striatum compared with some other brain regions, but detectable and functionally important. Immunohistochemistry experiments confirmed the efficiency and specificity of the conditional βarr2-KOs. In the NAc, βarr2 was detected in the majority of D2 cells in D1^βarr2-KO^ mice and D1 cells in D2^βarr2-KO^ mice. By contrast, βarr2 was largely absent in the majority of D1 cells in D1^βarr2-KO^ mice and D2 cells in D2^βarr2-KO^ mice, confirming a substantial cell type-specific reduction in βarr2 (Figure 1). Together, these results indicate cell type specificity of the KO and are consistent with previous work validating the conditional βarr2-KOs with immunohistochemistry and TRAP analysis (Urs, Gee, Pack et al., 2016).

While knocking out βarr2 had negligible effects on D1-MSN excitability in response to DA (Figure 2), eliminating βarr2 in D2-MSNs significantly altered DA-induced inhibition of these cells (Figure 3). The lack of DA-induced change in slope of F-I curves and half maximal current in NAc D2-MSNs of D2^βarr2-KO^ mice indicates that βarr2 normally contributes to DA-induced inhibition of D2-MSNs. To our knowledge, this is the first study to report the effects of endogenous βarr2 on multiple parameters of DA-induced changes in NAc D1- and D2-MSN excitability. UNC9994A, a βarr2-biased D2 partial agonist, recently was shown to be less effective than the D2/D3 agonist quinpirole at decreasing the excitability of D2-MSNs in the dorsal striatum (Urs, Gee, Pack et al., 2016). Thus, it appears that both G protein and βarr2 signaling are important for D2-induced inhibition of D2-MSNs.

The underlying mechanisms that lead to DA-induced changes in cellular excitability are complex. Some have reported that the D2/D3 agonist quinpirole has no effect on excitability (Lemos, Friend, Kaplan et al., 2016), while others have reported that quinpirole causes a decrease in intrinsic excitability (Gaval-Cruz, Goertz, Puttick et al., 2014; Urs, Gee, Pack et al., 2016). While the D2 cell response is largely dose-dependent, D1 agonists can facilitate either an increase or decrease in firing rate depending on the resting membrane potential (Hernández-López, Bargas, Surmeier et al., 1997). Because DA receptors are G-protein coupled receptors, they do not act directly on any specific ionic conductance but instead are modulators of multiple ionic conductances that either facilitate or inhibit intrinsic excitability. In these experimental conditions, we show that the endogenous agonist DA decreases excitability in D2-MSNs and increases excitability in D1-MSNs. The DA-induced decrease in D2-MSN excitability is partially dependent on βarr2 modulation, as we show that F-I slope is attenuated in D2^βarr2-KO^-MSNs. The exact biological mechanisms underlying how DA alters slope of the F-I curve is not completely understood. Because DA modulates sodium channel conductances (Surmeier & Kitai, 1993; Surmeier, Eberwine, Wilson et al., 1992), one possibility is that DA-dependent βarr2 signaling alters supra-threshold sodium conductances.

Whereas D1^βarr2-KO^ mice behaved similarly to controls in each locomotor test, D2^βarr2-KO^ mice exhibited dose-dependent reductions in cocaine- and morphine-induced locomotion (Figure 4) and blunted cocaine CPP (Figure 5). Similarly to D2^βarr2-KO^ mice, conventional βarr2-KO mice exhibited reduced locomotor responses to morphine (Bohn, Gainetdinov, Sotnikova et al., 2003); therefore, the present study provides cell-type specificity for βarr2’s attenuation of the locomotor-activating effects of morphine. Similarly to βarr2-KO mice, both D1^βarr2-KO^ and D2^βarr2-KO^ mice developed a CPP for a high dose of morphine that was not rewarding in controls; therefore, both D1 and D2 cells may encode a “ceiling” for the rewarding properties of morphine that is disrupted when βarr2 is eliminated. Unlike the conventional βarr2-KO mice (Bohn, Gainetdinov, Sotnikova et al., 2003), neither D1^βarr2-KO^ nor D2^βarr2-KO^ mice exhibited any alterations in baseline locomotor activity (Figure S2); therefore, that phenotype must be mediated by different cell types, and the drug-induced behavioral responses in our experiments are not confounded by changes in baseline activity.

Drug-induced DA release activates D2 receptors on D2-MSN, which promotes inhibition of these cells. D2-MSNs modulate the locomotor-activating effects of drugs of abuse, and both D1 and D2 receptors are involved in the rewarding properties of addictive drugs (Smith, Lobo, Spencer et al., 2013). Therefore, the electrophysiological finding that DA-induced inhibition of D2-MSNs is attenuated in D2^βarr2-KO^ mice is consistent with the behavioral finding of blunted cocaine-induced locomotion and reward. A recent calcium imaging study revealed that cocaine increases D1-MSN activity and decreases D2-MSN activity during CPP training, and inhibiting D1-MSNs prevents the development of CPP (Calipari, Bagot, Purushothaman et al., 2016). Considering the present data, these cocaine-induced changes in D1-MSNs are likely driven through G protein rather than βarr2 signaling, because D1^βarr2-KO^ mice displayed normal cocaine CPP. The current study shows that D2^βarr2-KO^ mice have attenuated cocaine CPP, and their D2-MSNs are less inhibited by DA. In light of the fact that cocaine reduces NAc D2-MSN activity (Calipari, Bagot, Purushothaman et al., 2016), the present data support the interpretation that βarr2 in D2-MSNs is necessary for maximal DA-induced inhibition of D2-MSNs in the presence of cocaine, and that this functionally contributes to the development of cocaine CPP.

Consistent with the present study, mice lacking βarr2 in all D2-containing cells or specifically in striatal D2 cells, but not in D1-containing cells, showed blunted amphetamine-induced locomotion (Urs, Gee, Pack et al., 2016). Viral vector-mediated striatal overexpression of D2 receptors with biased βarr2 signaling enhanced locomotor responses to amphetamine (Peterson, Pack, Wilkins et al., 2015), demonstrating that βarr2 in D2-MSNs can bidirectionally modulate psychostimulant-induced locomotion. The present data demonstrate that βarr2 in D2 cells modulates the locomotor-activating effects of cocaine in addition to amphetamine and demonstrate a novel role for βarr2 in D2 cells in mediating cocaine reward.

Mice completely lacking βarr2 originally were reported not to have deficits in cocaine-induced locomotion, locomotor sensitization to cocaine, or cocaine CPP (Bohn, Gainetdinov, Sotnikova et al., 2003). The discrepancy between these finding and the present study could be attributed to differences in the behavioral assays, doses of cocaine tested, or compensatory changes that may occur due to ubiquitous, lifelong loss of βarr2-KO. It is also possible that βarr2 facilitates cocaine responses in some cell types and opposes them in others. By eliminating βarr2 selectively in D2 cells, we may have uncovered a phenotype that is masked in the full KO. The present data revealing blunted cocaine responses in D2^βarr2-KO^ mice are consistent with recent findings showing that βarr2 is necessary for amphetamine-induced sensitization (Zurkovsky, Sedaghat, Ahmed et al., 2017), and D2^βarr2-KO^ mice have reduced acute responses to amphetamine (Urs, Gee, Pack et al., 2016).

In humans, decreased striatal D2 receptor availability is associated with cocaine abuse (Volkow, Fowler, Wang et al., 2009; Volkow, Fowler, Wang et al., 1993), whereas in the present study cocaine reward was blunted and D2 cellular responses to DA were impaired in D2^βarr2-KO^ mice. The differences between these findings is likely attributable to many factors. A reduction in βarr2 does not equate to a reduction in D2 receptor expression, and D2 receptor G protein signaling is intact in D2^βarr2-KO^ mice; it is not known how cocaine dependence might differentially affect striatal D2 G-protein versus βarr2 signaling. Finally, βarr2 knockout presumably affects many GPCRs in D2^βarr2-KO^ cells, in contrast with the relatively selective D2 receptor reduction in people suffering from cocaine abuse.

Because morphine-induced locomotion requires mesolimbic DA transmission (Hnasko, Sotak & Palmiter, 2005; Manzanedo, Aguilar & Minarro, 1999), the reduced ability of DA to inhibit D2-MSNs in D2^βarr2-KO^ mice could account for the attenuated locomotor response to morphine in these animals. Alternatively, the altered morphine responses of D2^βarr2-KO^ mice could be attributed to reduced βarr2 interactions with mu-opioid receptors on MSNs, rather than activation of DA receptors per se. Indeed, mu-opioid receptors are located on D2-MSNs, and deleting mu-opioid receptors only in forebrain GABAergic neurons (primarily MSNs) reduces heroin-induced locomotion yet enhances motivation to self-administer heroin (Charbogne, Gardon, Martín-García et al., 2017).

The possibility exists that βarr2 in DA neurons contributes to the observed behavioral effects, as βarr2 would be eliminated in D2 autoreceptor-containing neurons in D2^βarr2-KO^ mice. However, altered βarr2 in DA neurons is unlikely to explain the blunted behavioral responses to cocaine because cocaine-induced DA release is not altered in βarr2-KO mice (Bohn, Gainetdinov, Sotnikova et al., 2003), and mice with βarr2 eliminated only in D2-MSNs, not including DA neurons, similarly display attenuated amphetamine-induced locomotion (Urs, Gee, Pack et al., 2016). βarr2-KO mice have greater morphine-induced DA release in the NAc (Bohn, Gainetdinov, Sotnikova et al., 2003). While the morphine CPP of D2^βarr2-KO^ mice is consistent with increased morphine-induced DA release, this neural mechanism is unlikely to mediate the behavior. D1^βarr2-KO^ mice similarly showed a CPP for every tested dose of morphine, yet would not have altered levels of βarr2 in DA neurons because DA neurons do not contain D1 receptors (Weiner, Levey, Sunahara et al., 1991). The hippocampus is an alternative region that could contribute to the increased morphine CPP, as hippocampal D1 and D2 receptors can modulate morphine CPP (Assar, Mahmoudi, Farhoudian et al., 2016). The prefrontal cortex is functionally connected to the brain reward system and could also conceivably contribute. Indeed, a limitation of the transgenic conditional KOs is that βarr2 in all D1- and D2-expressing cells is targeted. While the DA-dependent behaviors tested here are largely mediated through striatal circuitry, and our results are consistent with viral vectors targeting only striatal D2 receptors (described below), we cannot rule out the possibility that βarr2 in D1 or D2 containing cells in other brain regions contribute to the observed behavioral effects.

βarr2 has multiple cellular functions including desensitizing and internalizing GPCRs, as well as initiating intracellular signaling independent of traditional G protein signaling (Smith & Rajagopal, 2016). While the present study demonstrates that βarr2 in D2 neurons modulates the locomotor-activating and rewarding properties of addictive drugs, the precise intracellular actions of βarr2 that mediate these effects are still being determined. βarr2 signaling complexes with JNK and phospho-ERK have been implicated in mediating the locomotor-effects of morphine (Mittal, Tan, Egbuta et al., 2012; Urs, Daigle & Caron, 2011), and GSK3β in D2-MSNs has been shown to modulate amphetamine-induced locomotion (Urs, Snyder, Jacobsen et al., 2012). Futures studies should determine the specific signaling mechanisms in D2 cells mediating the altered behavioral responses to cocaine and morphine revealed in the present study.

Experiments using functionally selective D2 receptors in indirect pathway MSNs that are biased for either βarr (primarily βarr2) or G protein signaling showed that either mechanism is sufficient to support cocaine-induced locomotion, although the G protein group appeared to have lower cocaine-induced activity than the βarr group (Rose, Pack, Peterson et al., 2018), consistent with the present data. Another group reported identical cocaine-induced locomotion in mice with either βarr- or G protein-selective D2 receptors (Donthamsetti, Gallo, Buck et al., 2018); however, they used a higher dose of cocaine which may overwhelm and mask βarr2-mediated changes in locomotion. Although both βarr and G protein signaling in D2-MSNs contribute to amphetamine-induced locomotion, neither independently drives the full effect (Rose, Pack, Peterson et al., 2018). Viral overexpression of wild-type or βarr-biased D2 receptors in D2-MSNs potentiates locomotor responses to amphetamine, while overexpression of G protein-biased D2 receptors is less effective. Together, these experiments reveal that βarr signaling in D2-MSNs functionally contributes to locomotor responses to psychostimulants.

The present study highlights the importance of testing multiple doses of drugs, as different neurobiological mechanisms may contribute to responses at different doses. For example, as discussed above, mice with D2 receptors biased for G protein signaling have slightly reduced locomotor activity to 10 mg/kg cocaine (Rose, Pack, Peterson et al., 2018), but experiments using higher doses of cocaine did not detect effects from βarr2 manipulations (Bohn, Gainetdinov, Sotnikova et al., 2003; Donthamsetti, Gallo, Buck et al., 2018), consistent with our results. Similarly, it is only with higher doses of morphine (20 mg/kg) where D2-derived βarr2 contributions to morphine-induced locomotion are observed. Together, these data suggest that both G protein and βarr signaling contribute to the locomotor-activating effects of psychostimulants and opioids. G protein signaling in D2 cells appears sufficient for driving locomotion in response to low and high doses of cocaine, while βarr2’s contributions to cocaine-induced locomotion is most notable at the moderate 10 mg/kg dose. G protein signaling is sufficient for morphine-induced locomotion as 5 and 10 mg/kg, but βarr2 in D2 cells is necessary for normal locomotor responses to the higher dose of 20 mg/kg morphine.

Furthermore, effective doses can vary by behavioral assay because locomotor-activating effects do not always correlate with the rewarding properties of drugs. Although D2^βarr2-KO^ mice did not exhibit altered locomotor responses to 5 mg/kg cocaine, they did not find this dose rewarding whereas controls did. At the 10 mg/kg dose of cocaine, D2^βarr2-KO^ mice showed reduced responses to both the rewarding and locomotor-activating effects of the drug, and at 20 mg/kg cocaine, these mice showed similar responses as controls in both assays. Additionally, whereas all three genotypes showed similar CPP and locomotor responses to 5 and 10 mg/kg morphine, D2^βarr2-KO^ mice had blunted locomotor but *enhanced* CPP responses to 20 mg/kg morphine. These findings indicate that both G protein and βarr2 signaling in D2 cells modulate morphine’s effects, and βarr2’s contributions are bidirectional at higher doses enhancing reward but attenuating locomotion. For behavioral pharmacology studies, we advocate for testing multiple drug doses in each type of assay when possible: different neurobiological mechanisms can contribute to different drug-induced behavioral effects in a dose-dependent manner.

The present study revealed that βarr2 in D2 cells can modulate the locomotor-activating and rewarding effects of cocaine and morphine. Because βarr2’s effects varied depending on the behavioral assay and drug, investigating βarr2’s involvement in the reinforcing properties of various drugs of abuse will be important. Overexpression of wild-type D2 receptors on indirect pathway MSNs has been shown to enhance operant responding for a food reward in a progressive ratio test; however, overexpression of βarr-biased D2 receptors did not (Donthamsetti, Gallo, Buck et al., 2018), indicating that both G protein and βarr signaling are important for the motivation to obtain rewards. Future studies should examine the role of βarr2 in specific cell types in mediating complex motivated behaviors and reinforcement for both natural and drug rewards.

## Acknowledgments

We would like to thank Dr. Jeff Benovic for generously supplying the anti-βarr2 primary antibody used in this study and Pamela Romero for technical assistance. This work was funded by NIH R01DA038453 to DW and CAP. KAPS was funded by NIH F32NS098615. AKP was funded by the Brown Foundation Fellowship. Development and characterization of the floxed βarrestin2 mouse line was supported by 5R3-MH-073853 to MGC. This research project was supported in part by the Emory University Integrated Cellular Imaging Microscopy Core.

MGC is an inventor on patents and patent applications pertaining to an adjunct 5-HTP SR method of treatment for treatment-resistant depression. MGC owns equity in Evecxia, a company aiming at developing a 5-HTP SR drug. MGC also owns stock in Acadia Pharmaceuticals and has received compensation in the form of honoraria for lecturing at various academic institutions. KAPS, AKP, SLK, LCL, NMU, CAP, and DW have nothing to disclose.

## Author contributions

DW and CAP designed the study. KAPS collected and analyzed the locomotor, CPP, and immunohistochemistry data, and SLK and KAPS collected and analyzed the locomotor sensitization data. AKP collected and analyzed the electrophysiology data. NMU and MGC generated the floxed βarr2 mouse. CL bred and genotyped the conditional βarr2-KO mice. KAPS drafted the manuscript and figures with critical revisions provided by AKP, DW, CP, SLK, NMU, and MGC. All authors reviewed the manuscript and approved the final version.

## References

Aragona BJ, Cleaveland NA, Stuber GD, Day JJ, Carelli RM, Wightman RM (2008) Preferential enhancement of dopamine transmission within the nucleus accumbens shell by cocaine is attributable to a direct increase in phasic dopamine release events. J Neurosci 28:8821–8831.

Assar N, Mahmoudi D, Farhoudian A, Farhadi MH, Fatahi Z, Haghparast A (2016) D1- and D2-like dopamine receptors in the CA1 region of the hippocampus are involved in the acquisition and reinstatement of morphine-induced conditioned place preference. Behav Brain Res 312:394–404.

Beaulieu J-M, Gainetdinov RR (2011) The physiology, signaling, and pharmacology of dopamine receptors. Pharmacological reviews 63:182–217.

Beaulieu JM, Sotnikova TD, Marion S, Lefkowitz RJ, Gainetdinov RR, Caron MG (2005) An Akt/beta-arrestin 2/PP2A signaling complex mediates dopaminergic neurotransmission and behavior. Cell 122:261–273.

Bohn LM, Gainetdinov RR, Sotnikova TD, Medvedev IO, Lefkowitz RJ, Dykstra LA, Caron MG (2003) Enhanced rewarding properties of morphine, but not cocaine, in βarrestin-2 knock-out mice. J Neurosci 23:10265–10273.

Calipari ES, Bagot RC, Purushothaman I, Davidson TJ, Yorgason JT, Peña CJ, Walker DM, Pirpinias ST, Guise KG, Ramakrishnan C (2016) In vivo imaging identifies temporal signature of D1 and D2 medium spiny neurons in cocaine reward. Proceedings of the National Academy of Sciences 113:2726–2731.

Charbogne P, Gardon O, Martín-García E, Keyworth HL, Matsui A, Mechling AE, Bienert T, Nasseef T, Robé A, Moquin L (2017) Mu opioid receptors in gamma-aminobutyric acidergic forebrain neurons moderate motivation for heroin and palatable food. Biological psychiatry 81:778–788.

Di Chiara G, Imperato A (1988) Drugs abused by humans preferentially increase synaptic dopamine concentrations in the mesolimbic system of freely moving rats. Proc Natl Acad Sci 85:5274–5278.

Donthamsetti P, Gallo EF, Buck DC, Stahl EL, Zhu Y, Lane JR, Bohn LM, Neve KA, Kellendonk C, Javitch JA (2018) Arrestin recruitment to dopamine D2 receptor mediates locomotion but not incentive motivation. Molecular Psychiatry:1.

Floresco SB (2015) The Nucleus Accumbens: An Interface Between Cognition, Emotion, and Action. In: Annual Review of Psychology, Vol 66. Fiske ST (ed). Annual Reviews: Palo Alto. pp 25–52.

Gaval-Cruz M, Goertz RB, Puttick DJ, Bowles DE, Meyer RC, Hall RA, Ko D, Paladini CA, Weinshenker D (2014) Chronic loss of noradrenergic tone produces β-arrestin2-mediated cocaine hypersensitivity and alters cellular D2 responses in the nucleus accumbens. Addiction biology.

Hernández-López S, Bargas J, Surmeier DJ, Reyes A, Galarraga E (1997) D1 receptor activation enhances evoked discharge in neostriatal medium spiny neurons by modulating an L-type Ca2+ conductance. Journal of Neuroscience 17:3334–3342.

Hikida T, Morita M, Macpherson T (2016) Neural mechanisms of the nucleus accumbens circuit in reward and aversive learning. Neuroscience research 108:1–5.

Hnasko TS, Sotak BN, Palmiter RD (2005) Morphine reward in dopamine-deficient mice. Nature 438:854–857.

Kupchik YM, Brown RM, Heinsbroek JA, Lobo MK, Schwartz DJ, Kalivas PW (2015) Coding the direct/indirect pathways by D1 and D2 receptors is not valid for accumbens projections. Nature neuroscience 18:1230.

Lemos JC, Friend DM, Kaplan AR, Shin JH, Rubinstein M, Kravitz AV, Alvarez VA (2016) Enhanced GABA Transmission Drives Bradykinesia Following Loss of Dopamine D2 Receptor Signaling. Neuron 90:824–838.

Manzanedo C, Aguilar M, Minarro J (1999) The effects of dopamine D2 and D3 antagonists on spontaneous motor activity and morphine-induced hyperactivity in male mice. Psychopharmacology 143:82–88.

Mittal N, Tan M, Egbuta O, Desai N, Crawford C, Xie C-W, Evans C, Walwyn W (2012) Evidence that behavioral phenotypes of morphine in β-arr2−/− mice are due to the unmasking of JNK signaling. Neuropsychopharmacology 37:1953.

Pardo-Garcia TR, Garcia-Keller C, Penaloza T, Richie CT, Pickel J, Hope BT, Harvey BK, Kalivas PW, Heinsbroek JA (2019) Ventral pallidum is the primary target for accumbens D1 projections driving cocaine seeking. Journal of Neuroscience 39:2041–2051.

Peterson SM, Pack TF, Wilkins AD, Urs NM, Urban DJ, Bass CE, Lichtarge O, Caron MG (2015) Elucidation of G-protein and beta-arrestin functional selectivity at the dopamine D2 receptor. Proc Natl Acad Sci U S A 112:7097–7102.

Pierce RC, Kumaresan V (2006) The mesolimbic dopamine system: The final common pathway for the reinforcing effect of drugs of abuse? Neurosci Biobehav Rev 30:215–238.

Porter-Stransky KA, Centanni SW, Karne SL, Odil LM, Fekir S, Wong JC, Jerome C, Mitchell HA, Escayg A, Pederson NP (2019) Noradrenergic transmission at alpha1-adrenergic receptors in the ventral periaqueductal gray modulates arousal. Biological Psychiatry 85:237–247.

Porter-Stransky KA, Seiler JL, Day JJ, Aragona BJ (2013) Development of behavioral preferences for the optimal choice following unexpected reward omission is mediated by a reduction of D 2-like receptor tone in the nucleus accumbens. European Journal of Neuroscience 38:2572–2588.

Porter-Stransky KA, Weinshenker D (2017) Arresting the Development of Addiction: The Role of beta-Arrestin 2 in Drug Abuse. J Pharmacol Exp Ther 361:341–348.

Rose SJ, Pack TF, Peterson SM, Payne K, Borrelli E, Caron MG (2018) Engineered D2R variants reveal the balanced and biased contributions of G-protein and β-arrestin to dopamine-dependent functions. Neuropsychopharmacology 43:1164.

Schank JR, Ventura R, Puglisi-Allegra S, Alcaro A, Cole CD, Liles LC, Seeman P, Weinshenker D (2006) Dopamine [beta]-hydroxylase knockout mice have alterations in dopamine signaling and are hypersensitive to cocaine. Neuropsychopharmacology 31:2221.

Smith JS, Rajagopal S (2016) The β-arrestins: multifunctional regulators of G protein-coupled receptors. J Biol Chem 291:8969–8977.

Smith RJ, Lobo MK, Spencer S, Kalivas PW (2013) Cocaine-induced adaptations in D1 and D2 accumbens projection neurons (a dichotomy not necessarily synonymous with direct and indirect pathways). Current Opinion in Neurobiology 23:546–552.

Surmeier D, Kitai S (1993) D1 and D2 dopamine receptor modulation of sodium and potassium currents in rat neostriatal neurons. In: Progress in Brain Research. Elsevier. pp 309–324.

Surmeier DJ, Eberwine J, Wilson CJ, Cao Y, Stefani A, Kitai ST (1992) Dopamine receptor subtypes colocalize in rat striatonigral neurons. Proc Natl Acad Sci U S A 89:10178–10182.

Urs NM, Daigle TL, Caron MG (2011) A dopamine D1 receptor-dependent beta-arrestin signaling complex potentially regulates morphine-induced psychomotor activation but not reward in mice. Neuropsychopharmacology 36:551–558.

Urs NM, Gee SM, Pack TF, McCorvy JD, Evron T, Snyder JC, Yang X, Rodriguiz RM, Borrelli E, Wetsel WC, Jin J, Roth BL, O’Donnell P, Caron MG (2016) Distinct cortical and striatal actions of a β-arrestin–biased dopamine D2 receptor ligand reveal unique antipsychotic-like properties. Proc Natl Acad Sci 113:E8178–E8186.

Urs NM, Snyder JC, Jacobsen JP, Peterson SM, Caron MG (2012) Deletion of GSK3β in D2R-expressing neurons reveals distinct roles for β-arrestin signaling in antipsychotic and lithium action. Proceedings of the National Academy of Sciences 109:20732–20737.

Vander Weele CM, Porter-Stransky KA, Mabrouk OS, Lovic V, Singer BF, Kennedy RT, Aragona BJ (2014) Rapid dopamine transmission within the nucleus accumbens: Dramatic difference between morphine and oxycodone delivery. Eur J Neurosci 40:3041–3054.

Volkow N, Fowler J, Wang G, Baler R, Telang F (2009) Imaging dopamine’s role in drug abuse and addiction. Neuropharmacology 56:3–8.

Volkow ND, Fowler JS, Wang GJ, Hitzemann R, Logan J, Schlyer DJ, Dewey SL, Wolf AP (1993) Decreased dopamine D2 receptor availability is associated with reduced frontal metabolism in cocaine abusers. Synapse 14:169–177.

Weiner DM, Levey AI, Sunahara RK, Niznik HB, O’Dowd BF, Seeman P, Brann MR (1991) D1 and D2 dopamine receptor mRNA in rat brain. Proceedings of the National Academy of Sciences 88:1859–1863.

Zurkovsky L, Sedaghat K, Ahmed MR, Gurevich VV, Gurevich EV (2017) Arrestin-2 and arrestin-3 differentially modulate locomotor responses and sensitization to amphetamine. Neuropharmacology 121:20–29.

