## Supplemental Information for "Loss of β-arrestin2 in D2 cells alters neuronal excitability in the nucleus accumbens and behavioral responses to psychostimulants and opioids"

**Supplementary Materials and Methods**

**Immunohistochemistry.** To confirm that βarr2 was reduced in D1- and D2-MSNs of D1^βarr2-KO^ and D2^βarr2-KO^ mice, 40 µm thick striatal sections were obtained and processed as previously described (Porter-Stransky, Centanni, Karne et al., 2019). Anti-βarr2 primary antibody (generated in rabbit, generously gifted from Jeff Benovic, Thomas Jefferson University) was used at 1:300 concentration. Because βarr2 levels are relatively low in the striatum, Tyramide Signal Amplification (Alexa Flour 488 Tyramide SuperBoost Kit, goat anti-rabbit IgG, Thermo Fisher) was used to augment βarr2 detection. To control for any non-specific binding of the secondary antibody and the Tyramide Signal Amplification, a subset of tissue was processed as described above except that the primary antibody was omitted. Endogenous fluorescence from the tdTomato transgene was used to visualize D1 or D2 cells in each line. Images were captured at 40x on an Olympus FV1000 confocal microscope using Olympus FluoView software. Image J was used to quantify the percentage of cells containing DAPI, tdTomato, and βarr2 immunofluorescence.

**Slice electrophysiology.** NAc neurons identified in D1^βarr2-KO/tdTomato^ and D2^βarr2-KO/tdTomato^ mice, or D1-tdTomato and D2-GFP as control mice, were used for electrophysiological recordings. Both males and females 8-12 weeks of age were used. Mice were anesthetized with a lethal dose of isoflurane and decapitated. The brains were quickly removed and placed into oxygenated cutting aCSF at 37⁰ C containing (in mM): 126 NaCl, 2.5 KCl, 1.25 NaH_2_PO4, 2 MgCl_2_, 2 CaCl_2_, 10 dextrose, 25 NaHCO_3_, 1.3 ascorbic acid, 2.4 sodium pyruvate and 0.05 glutathione, along with 2 μM kynurenic acid. Coronal brain slices containing the NAc (250 μm) were cut using a vibrating tissue slicer (Microm HM 650V, Thermo Fisher Scientific, Waltham, MA). The slices were then transferred to an incubation chamber containing the same aCSF without kynurenic acid, but with 10 µM MK-801 (NMDA receptor antagonist) for 30 minutes prior to recordings. Slices were then taken out of the incubation chamber and stored at room temperature in aCSF containing no kynurenic acid or MK-801 for the remainder of the day. The slices were transferred to a recording chamber for the experiments, where they were submerged in oxygenated aCSF equilibrated with 95%O_2_-5%CO_2_ at a pH of 7.2, containing (in mM): 126 NaCl, 2.5 KCl, 1.25 NaH_2_PO_4_, 2 MgCl_2_, 2 CaCl_2_, 10 dextrose, 25 NaHCO_3_, and 2.4 sodium pyruvate. The slices were superfused with 34 – 36°C aCSF at a rate of 2 ml/minute. The cells were visualized using gradient contrast illumination through a 60X water-immersion lens attached to an Olympus BX51 upright microscope (Olympus, Center Valley, PA). Patch pipettes were pulled from borosilicate glass (o.d. 1.5 mm, i.d. 0.84 mm) using a P-1000 Flaming/Brown electrode puller (Sutter Instruments, Novato, CA). Pipettes were filled with a solution containing (in mM): 138 K-gluconate, 10 HEPES, 0.0001 CaCl_2_, 0.2 ethylene glycol tetraacetic acid, 4 NaATP, 0.4 NaGTP and 2 MgCl_2_, with an osmolarity of 270–275 mOsm and adjusted to a pH of 7.3 with potassium hydroxylase (KOH). Recordings were made using a MultiClamp 700B amplifier (Molecular Devices, Sunnyvale, CA). Signals were digitized at 30-50 kHz and saved to a hard drive for analysis using the software program AxoGraph X (AxoGraph Scientific, Berkeley, CA). D1-MSNs or D2-MSNs were identified using epifluorescence microscopy to identify somatic tdTomato or EGFP expression to confirm cell identity before breaking into the cell. Physiological identification of MSNs in the NAc core included the following: hyperpolarized membrane potential (< −70 mV), low input resistance (< 350 MΩ), and delayed spiking upon depolarizing current injection. DA (60 μM) was applied to the slice by superfusion. All experiments were performed in the presence of 5 μM NBQX (AMPA receptor antagonist), 10 μM MK-801 (NMDA receptor antagonist), 100 μM picrotoxin (GABA_A_ receptor antagonist), 10 μM mecamylamine (nicotinic receptor antagonist), 1 μM atropine (muscarinic receptor antagonist), and 50 nM CGP 54626 (GABA_B_ receptor antagonist). All drugs were obtained from Tocris Bioscience or Sigma-Aldrich. In current-clamp configuration, current was injected for 1 sec at 25-pA step intervals (-50 – 500 pA) with 5 sec between each pulse. An input/output curve was obtained under baseline conditions before and after superfusing 10 ml of a 60 μM solution of DA HCl for approximately 5 minutes (Manvich, Petko, Branco et al., 2019; Planert, Berger & Silberberg, 2013). Using the identical preparation, we have shown that sulpiride reverses 60 μM DA-induced decreases in D2-MSN firing, demonstrating selectivity of this concentration of DA for DA receptors. Stock solutions of DA HCl were made fresh daily.

Input resistance was calculated by taking the average membrane potential at steady state during specific current injection amplitude levels (-50 – 75 pA). Slope values for each F-I curve were determined using linear fit through the linear, or primary region of the F-I curve, which was assessed individually for each cell and defined by firing rates at the first four consecutive current injections above rheobase. Rheobase was measured at the point where the linear fit line crossed the x-axis and was measured within each individual cell’s F-I curve. Maximum firing rate was measured by taking each cell’s maximum firing frequency right before the cell underwent depolarization block.

**Drug-induced locomotion.** San Diego Instruments (La Jolla, CA) locomotor chambers were used for locomotor testing. Cages (22 x 43 x 22 cm) with corncob bedding were situated within a 25 x 47 cm stainless steel frame that passed 7 infrared beams through the long wall of the locomotor activity chamber at 5.5 cm intervals. Mice underwent three 1-h habituation sessions in the chambers and subsequently received one drug-induced locomotor test per week. Drug order was counterbalanced across subjects. Each test session began with a 90-min habituation period followed by administration of saline, cocaine, or morphine (5, 10, or 20 mg/kg, i.p.), and drug-induced locomotion was monitored for 2 h. The primary outcome measure was ambulations, defined as consecutive photo-beam breaks, collected in consecutive 5-min bins.

To test whether baseline locomotor activity of controls bred from D1-Cre x floxed βarr2 mice and D2-Cre x floxed βarr2 mice differed, saline-induced locomotion for 30-minutes post injection was quantified between controls (homozygous floxed βarr2 mice) with a D1-Cre vs D2-Cre parent.

To further test the role of βarr2 in D2 cell in modulating cocaine-induced locomotion, the locomotor activity of control and D2^βarr2-KO^ mice was recorded in the CPP chambers during the 30-minute conditioning sessions to 10 mg/kg cocaine.

**Locomotor Sensitization.** Following a 60-min habituation period, mice received cocaine (10 mg/kg, i.p.), and locomotor activity was measured for 60 min. This procedure occurred daily for 7 consecutive days.

**CPP.** On days 1 and 5, mice were placed in the center chamber midday and allowed to freely explore all chambers for 20 min. On days 2-4, mice completed two 30-min sessions per day: in the morning mice received a saline injection and were confined to one side of the apparatus, and in the afternoon mice received an injection of saline or drug (5, 10, or 20 mg/kg, i.p. of cocaine or morphine) and were confined to the other side of the apparatus. A preference score was calculated on days 1 (pre-test) and 5 (post-test) by subtracting time spent in the saline-paired compartment from time spent in the drug-paired compartment. For preference score calculations of subjects who received saline in both compartments, the “saline” side was the side in which paired with the saline injection in the morning session and the “drug” side was the side in which saline was paired in the afternoon session.

**Supplementary Figures**

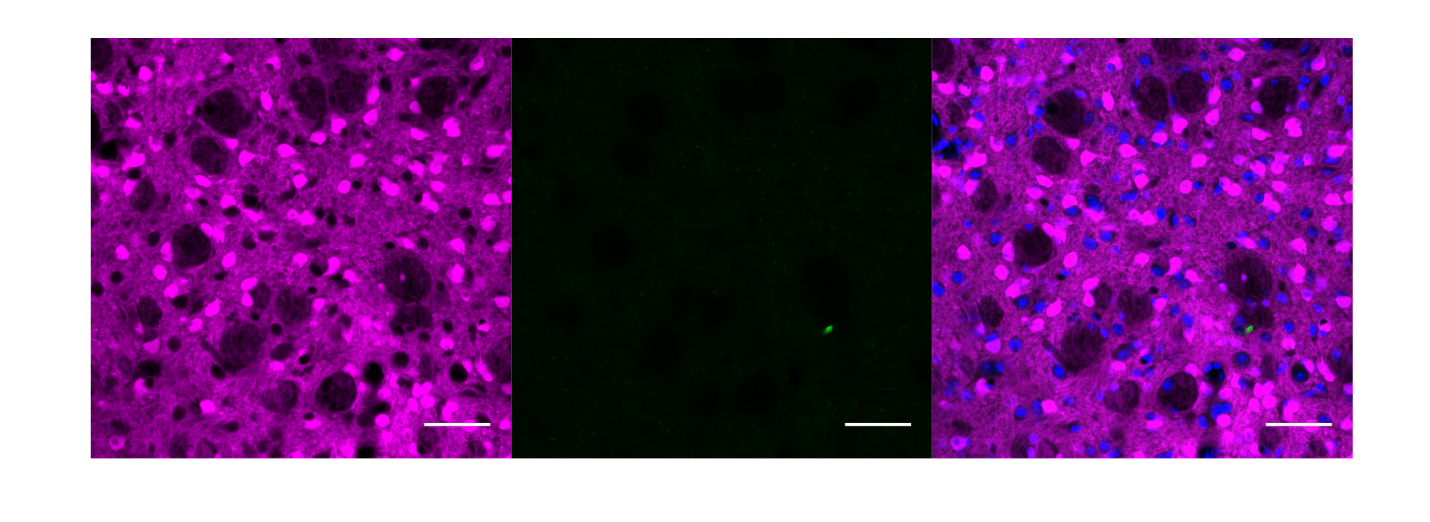

**Figure S1.** **Immunohistochemistry secondary antibody control.** To determine whether non-specific staining from the Tyramide Signal Amplification secondary antibody kit was occurring and impacting the observed βarr2 signal, striatal sections from a D1^βarr2-KO/tdTomato^ mouse were processed identically as the tissue in Figure 1 except that the anti-βarr2 primary antibody was omitted. Endogenous tdTomato fluorescence (magenta) indicates D1-MSNs. The minimal green staining (center pane), demonstrates that the green signal in Figure 1 was not due to non-specific binding of the secondary antibody. Blue in the merged images represents DAPI. All scale bars = 50 µm.

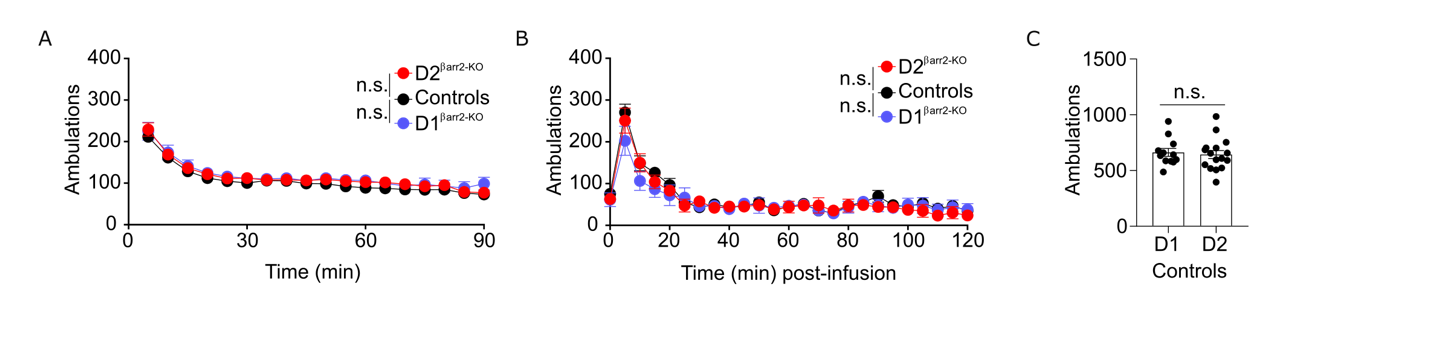

**Figure S2. Elimination of βarr2 in D1 or D2 cells has no effect on baseline or saline-induced locomotion. A-B)** D1^βarr2-KO^ and D2^βarr2-KO^ mice had similar locomotor responses to a novel environment (A) and a saline injection (B) as controls (n = 11 mice per genotype). **C)** Saline-induced locomotion did not differ between controls bred from D1-Cre x floxed βarr2 mice (D1; n = 12) and controls bred from D2-Cre x floxed βarr2 mice (D2; n = 16). n.s. = not statistically significant, error bars represent SEM.

**Supplementary References**

Manvich DF, Petko AK, Branco RC, Foster SL, Porter-Stransky KA, Stout KA, Newman AH, Miller GW, Paladini CA, Weinshenker D (2019) Selective D2 and D3 receptor antagonists oppositely modulate cocaine responses in mice via distinct postsynaptic mechanisms in nucleus accumbens. *Neuropsychopharmacology*.

Planert H, Berger TK, Silberberg G (2013) Membrane properties of striatal direct and indirect pathway neurons in mouse and rat slices and their modulation by dopamine. *PloS one* 8:e57054.

Porter-Stransky KA, Centanni SW, Karne SL, Odil LM, Fekir S, Wong JC, Jerome C, Mitchell HA, Escayg A, Pederson NP (2019) Noradrenergic transmission at alpha1-adrenergic receptors in the ventral periaqueductal gray modulates arousal. *Biological Psychiatry* 85:237-247.
